# Acetylcholine as a trigger of the somatic exocytosis in Retzius neurons

**DOI:** 10.1101/2020.12.01.405951

**Authors:** Tatiana A. Kazakova, Oleg N. Suchalko, Alexey D. Ivanov, Anna V. Alova, George V. Maksimov

## Abstract

The redistribution of vesicles containing serotonin in leech neurons was studied using the fluorescent, scanning ion-conductance, and laser phase microscopy methods. During acetylcholine receptor (AChR) activation in Retzius neurons, the changes of Ca^2+^ desorption, cellular stiffness and the cell optical phase difference (OPD) were established. It was found that the amplitude of OPD changes in the near-membrane area (membrane and near-membrane of the cytoplasm layers) increases upon AChR activation and this is, possibly, associated with the neurons vesicle redistribution. The decrease in the cell stiffness upon AChR activation suggests the crucial role of cytoskeleton for vesicle transport and release. Ca^2+^ rise in the cytoplasm during AChR activation may regulate the mitochondrial recruitment to regions with high energy demand for vesicle trafficking.

## 1. Introduction

The neurilemma depolarization of neuron plasma membrane is accompanied with the generation of action potential (AP) and increase in Na^+^- and Ca^2+^-influx into the cell. The neuron AP generation with different frequencies (rhythmic activity, RA) is based on an activation of the voltage-gated and ligand-operated channels and the long lasting neuron plasmalemma changes (surface membrane potential and membrane viscosity, reorganization of cytoplasm structures). The spontaneous electrical activity of neurons in leech ganglia can be modified by the certain neurotransmitters [1-3]. However, the regulatory mechanisms of RA modulation in neurons are still poorly understood.

The Retzius cells in the leech (Rz-neurons) produce and release serotonin (5-HT) that plays an important neuromodulator role and may regulate the RA [4-6]. Serotonin may act in three different ways in the nervous system: as a transmitter at the synapses; as a paracrine modulator upon diffusion at a distance from its release sites, and as a hormonal modulator by circulating in the blood stream [7]. The three modes can affect a single neuronal circuit. The secretion of serotonin can occur synaptically or extrasynaptically. The modulation of neural circuits requires the large amounts of signalling molecules that are released not only in the synapses but also from extrasynaptic sites in the soma, dendrites and axons. This secretion maintains transmitter concentrations in the extracellular spaces of not only near, but also distant neurons, glial cells and blood vessels [8]. The volume transmission in response to transmitter release from extrasynaptic sites, in the soma, dendrites and axons recently attracted more attention and was termed “extrasynaptic communication” [9]. Serotonin is stored in dense core vesicles of the leech Retzius cells. Electrical stimulation of the cells or a long-lasting increase of intracellular Ca^2+^ may trigger the serotonin exocytosis [10]. The serotonin release from the Retzius cells depends on the membrane potential shifts, the RA frequency, on L-type calcium channel activation and on calcium-induced calcium release [7]. However, there are no direct evidence how other neurotransmitters of the extracellular “broth” may modulate the serotonin exocytosis. What are the natural intrinsic stimuli that modify neuronal rhythmic activity to trigger the extrasynaptic release of serotonin?

In the leech ganglion there is a basal level of acetylcholine (ACh) presence that is counteracted by endogenous acetylcholinesterase (AChE) activity to regulate the membrane potential and support a sustained firing rate of Retzius neurons [11]. We propose that the basal ACh release may influence the output of these neuromodulatory serotonergic neurons.

A complex response to acetylcholine (ACh) is displayed by the Retzius cells bearing receptors with different nicotinic pharmacological profiles. The activation of nicotinic acetylcholine receptors (AChR) can cause the membrane potential changes and modification of Retzius cells RA [2,12-14]. But there is no information in the literature that ACh action can evoke the extrasynaptic exocytosis of serotonin in the Retzius cells.

There are scarce references about the interplay between acetylcholine and serotonin action on Retzius cells. The extracellular acetylcholine application to the bath exerts a short-term increase in the frequency of the spontaneous impulse activity (RA). On the contrary, presence of serotonin in the extracellular solution leads to a temporary deceleration of the impulse activity [2,13].

As revealed with electron microscopy significant ultrastructure changes occur during AchR action on Retzius cells: the increase of the number of electron-transparent neurosecretory granules, the swelling of the mitochondria and the endoplasmic reticulum, partial elimination of ribosomes [9]. But in case of serotonin application the neuron ultrastructure can hardly be distinguished from the normal. Investigation of the responses mediated by nicotinic receptors in well-characterized serotonergic neurons *in situ* is likely to provide more direct information on the physiological role of AChR activation in the regulation of RA and extrasynaptic communication.

In this study, we address the functional role(s) of the nicotinic receptors in the central nervous system of invertebrates in connection to extrasynaptic release of transmitter substances. For the first time we revealed that acetylcholine can cause *serotonin release in Retzius cells.* The 5-HT exocytosis molecular mechanisms during AChR activation in Retzius cell were investigated: membrane Ca^2+^ - desorption, cytoskeleton rearrangements (cellular stiffness) and the neuron cytoplasm optic phase profile (OPD)) dynamics. We propose that acetylcholine and serotonin present in the ganglion may cooperate to sustain the proper functioning of neuronal circuits.

## 2. Experimental Section

### 2.1. Cell Culture

Isolated Retzius cells from the leech segmented ganglia (*Herudo medicinalis*) were used. Isolated neurons were incubated in a special chamber in the solutions with or without acetylcoline (115 mM NaCl, 4mM KCl, 1mM CaCl_2_, 1 mM MgCl_2_, pH 7.4 at 18 °C and 10^−8^ M acetylcholine [16].

### 2.1. The imaging of neurons with dynamic phase microscopy

The imaging of isolated Retzius neurons with dynamic phase microscopy (DPM) was performed with a phase microscope Eiriscan. In contrast to traditional optical microscopy, based on recording the distribution of light intensity, the DPM makes it possible to determine directly the optical phase difference (OPD) distribution in an interference image. The optical system of DPM consists of a modified Linnik interferometer with a coherent light source (helium - neon laser, λ=633.3nm, 1.5 mWt) and a dissector image tube acting as a coordinate-sensitive photo detector [17] The parameters of OPD measurements were: a size of the view field 128×128 p (20 × 20 μm), time of measurement t = 14.7 s. A line for scanning of length 40 p (6.1 microns) was chosen across a cell in its various areas and periodic measurements of OPD phase heights were performed there. The obtained data were converted in ASCII-codes as two - dimension of rectangular matrixes by the size 40 × 400 and processed by Fourier transform in a standard MATLAB package. The Fourier transform allows to reveal the dominant frequency components from matrix of phase values. The control measurements were carried out on a silicon substrate in the same conditions near to a cell.

### 2.3. The imaging of neurons with fluorescent microscopy

The fluorescent microscope (Lumum I-3, LOMO, Russia) was used to obtain fluorescent data. A halogen lamp KGM 12-100 served as a source of light. To excite the fluorescence of CTC (10^−4^M), Rh123 (10^−5^M rodamin) and neuron autofluorescence, a combination of FS-1-6 and SZS 21-2 filters was used. The signal registration was carried out with a photometric nozzle of PMEL-1A using interference optical filters (transmission maximum in the region of 520 nm). The neurons were incubated during 10-15 minutes in the standard solution containing 10^−4^ M CTC or incubation with 10^−5^ M rodamin (Rh123) during 10-15 minutes and subsequent triple washing [16].

### 2.4. *The imaging of neurons with* Scanning ion conductance microscopy

The scanning ion conductance microscope is based on inverted optical microscope Eclipse Ti-2 (Nikon, Japan), that is placed on vibration isolation table STable (Supertech instruments, Hungary). The microscope contains several functional parts, such as scanning platform *Mechanical Stand* (ICAPPIC, UK), positioning system *Piezo control system* (ICAPPIC, UK), feedback control system *Universal Controller* (ICAPPIC, UK), and I/O interface Multiclamp 700B и Axon Digidata 1550B (Axon Instruments, USA). The stiffness evaluation was performed using Clarke’s method [18].

### 2.8. Statistics

The results were statistically processed using the GraphPad Prism software, version 8.02 (GraphPad Software, La Jolla California USA). During the study, 12 animals were used, in each of which Retzius cells was assessed. All data were normally distributed (according to the generalized D’Agostino-Pearson test for normal distribution, p <0.05).

## 3. Results

### 3.1. Retzius cell optic phase difference

The OPD of the Retzius cell is determined the neuron cytoplasm optical density and volume. The optic phase difference correlates with the neuron membrane or/and intracellular processes: (1). this phase shift may be caused by the processes that occur in the cytoplasm, such as cyclosis, exo- and endocytosis. The transport of different vesicles, volume changes of mitochondria and nucleus influence the optical density changes; (2). the optical density changes correlate with the near membrane volume and membrane fluidity. In these experiments, a distribution of OPDs of the Retzius cell was investigated (Fig.1A). The optical heterogeneity of neuron topography is associated with the different cytoplasm subcellular organelles and vesicles. In addition, the OPDs changes were found both in the cell and in the near-membrane region (a resolution of 10 nm) (Fig. 1B). It has been established that in control conditions (without AChR activation) cytoplasm of Retzius neurons had the small OPD changes in the perinuclear and the neurolemma regions (Fig.1B). Probably, the OPD changes in the near-membrane region are associated with the vesicle redistribution during Retzius neurons spontaneous RA [9].

**Figure. 1.**
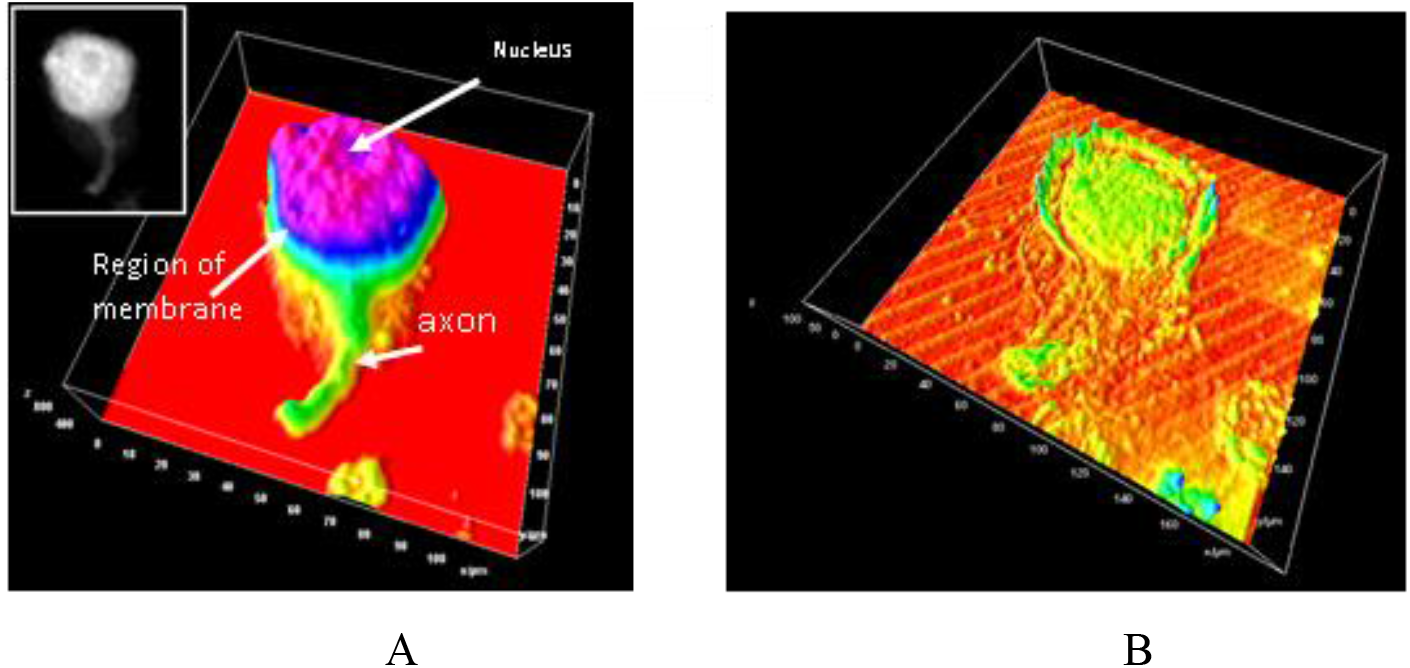
A - Retzius cell OPDs distributions (on the inset - the neuron light image). The arrows indicate the nucleus, the axon and the area corresponding to the cell membrane and the near membrane region; B - The distribution of the standard deviation of the cells OPDs;

### 3.2. Retzius cell optic phase difference during the AChR activation

During the Retzius cell AChR activation, the OPD amplitude and distribution changed: in the near nucleus cytoplasm region the OPD amplitude was increased, but in the near-membrane area (membrane and near-membrane of the cytoplasm layers) the OPD amplitude was decreased. Obviously, the neuron OPD changes observed during the AChR activation are due to the vesicles redistribution and changes in the refractive indices (n_i_) of the cytoplasm: 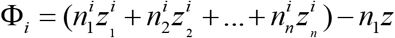 or the distance through which the light passes in the object (z_i_). The LPM resolution does not allow one to estimate the level of molecules changes and changes in the cytoplasm composition are also unlikely to make a significant contribution to OPD changes. It can be assumed that the main contribution to the OPD changes are connected with redistribution of cytoplasm intracellular vesicles. Using the OPD standard deviation data, the effect of AChR activation on changes in the amplitude and distribution of the maximum OPD in the cell near-membrane region were founded (Fig. 2A).

**Fig. 2.**
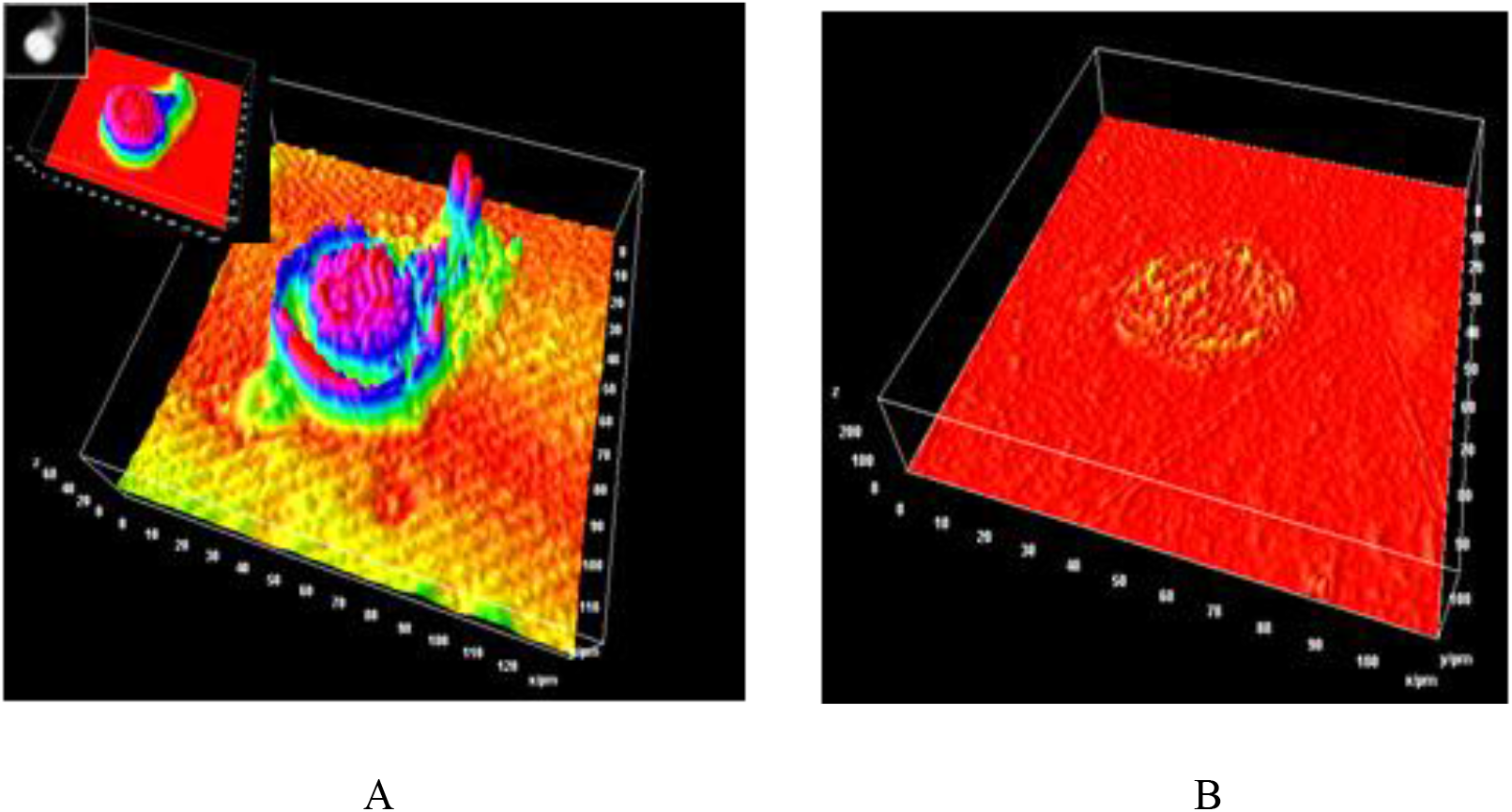
**A**. The standard deviation of the Retzius cell OPD changes during the AChR activation (shows areas with the most active changes in the cell phase profile). In the inserts, the same neuron OPD profile in transmitted light. **B** – the standard deviation of OPD changes in the different regions of the non-serotoninergic neuron.

There are two clusters of serotonin vesicles in the Retzius cell: the membrane and the perinuclear, located close to the Golgi apparatus. During neuron electrostimulation the release of near-membrane vesicles occurs in the first place; further stimulation causes the release of perinuclear vesicles [19]. It is likely that the AChR activation initiated both, but primary the near membrane vesicles movement. In addition, the maximal OPD changes in the nucleus region might be associated with the mitochondria redistribution, which are localized around nucleus or its volume changes. Note, in the non-serotoninergic neuron the ACh action did not lead to significant OPD changes (Fig.2B). So, the neuron OPD changes during the AChR activation are, probably, associated with the vesicles’ movement and subsequent serotonin exocytosis.

### 3.3. Retzius cellular stiffness during the AChR activation

The inspection of cellular stiffness was performed by means of scanning ion conductance microscopy. This approach provides information about cytoskeleton structure and its rearrangements [18]. During AChR activation in Retzius neurons the cell stiffness decreased twice after several minutes (Fig.3). The observed effect was reversible and five minutes later the treated cells returned to their initial state. This suggests that the near-membrane cytoskeleton reorganization could facilitate the arrival of vesicles from the neuron soma and promote prolonged exocytosis of serotonin. Effects of AChR activation on cell stiffness after consistent application of Glu and ACh neurotransmitters in the Retzius neurons. Both experiments show that the application of Glu was insufficient to cause any significant changes in cell stiffness due to low Glu-receptor density in leech neurons. At the same time, the application of ACh leads to a decrease in cell stiffness, which restores in a short span of time. The repeated application causes the same effect. Perhaps, the effect may be explained by AChR activation and exocytosis of serotonin-containing vesicles.

**Fig 3.**
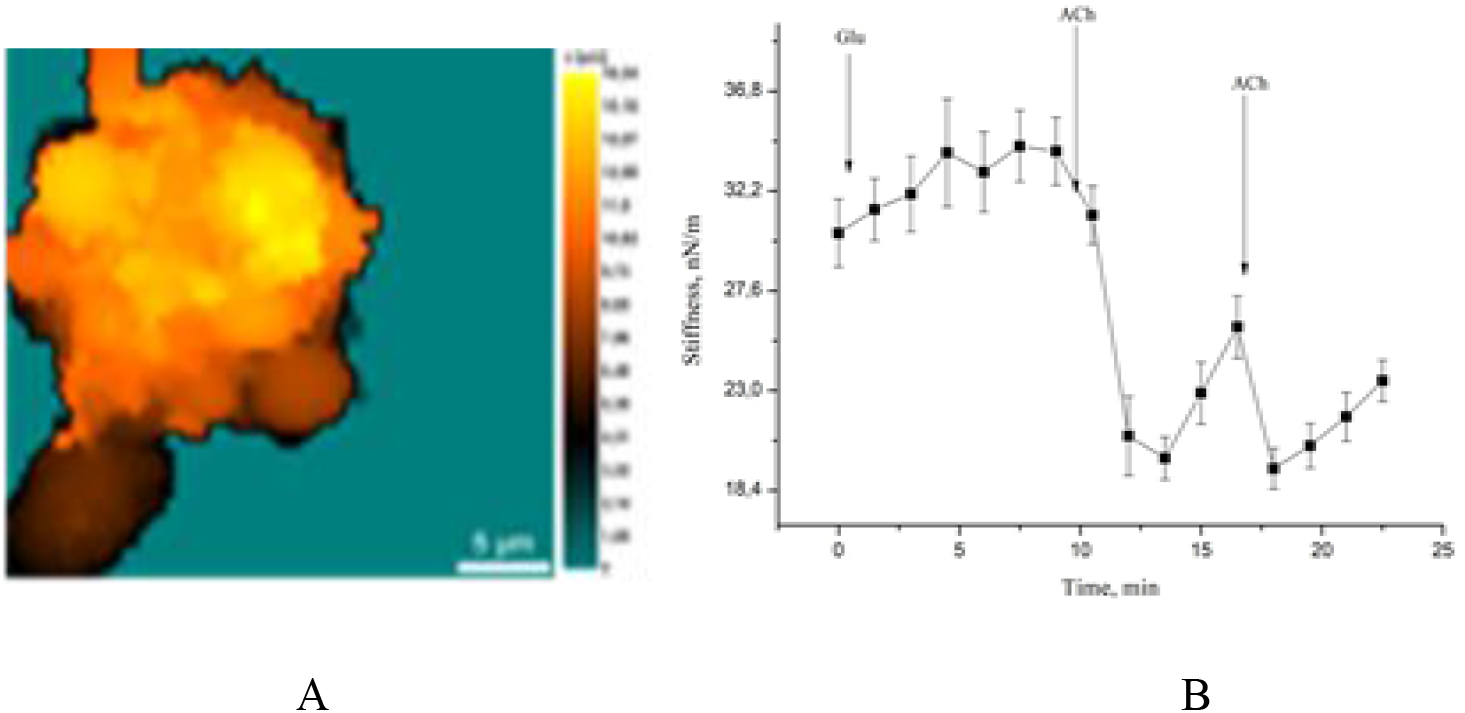
Retzius cellular stiffness distribution (A) and effect of AChR activation in the neurons stiffness (B).

### 3.4. Retzius cell membrane-bound calcium and the mitochondria inner membrane potential shift during the AChR activation

The neuron plasma membrane proteins and the mobility of cytoskeleton proteins are important factor for the vesicle redistribution during AchR activation [19-20]. It was found, that ACh receptor activation (not serotonin action) can change the level of membrane-bound calcium (Fig. 4). The amount of the bound Ca^2+^ was estimated by fluorescent microscopy with a fluorescent dye chlortetracycline (CTC). The decrease in the intensity of CTC fluorescence corresponds to the decrease in the amount of bound Ca^2+^.The AChR stimulation caused that the fraction of membrane-bound Ca^2+^ started to decrease after several minutes. This leads to increase in calcium concentration in the cytoplasm. The Ca^2+^ desorption may not only change the cell excitation threshold but facilitate the process of vesicles exocytosis.

**Fig. 4.**
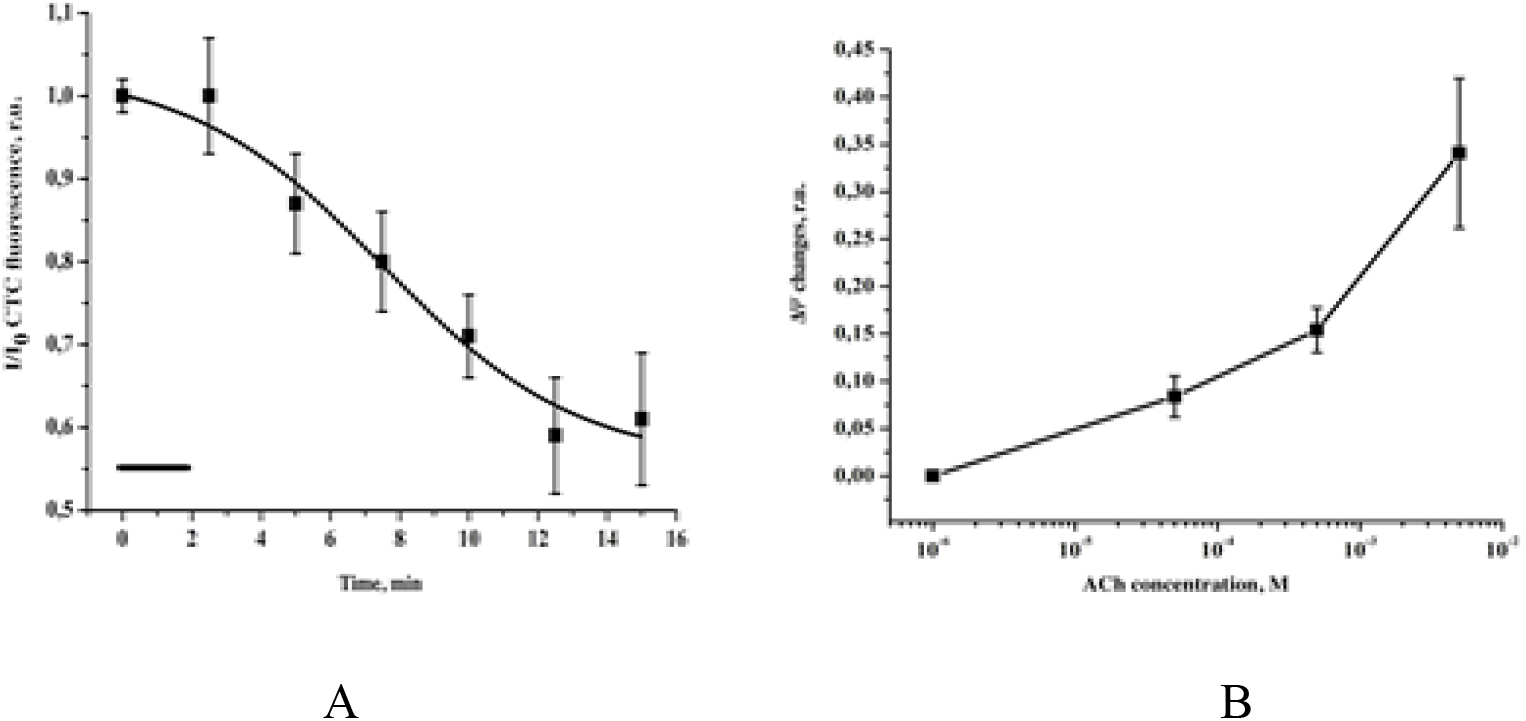
**A** The Retzius neuron membrane-bound calcium during the Retzius neuron AChR activation **B**. The mitochondria inner membrane potential shift in Retzius neurons as a function of Ach concentration.

In the next experiments, the mitochondrial membrane potential (ΔΨ) changes in Retzius cells during AChR activation were studied. We assessed mitochondrial membrane potential using the Rh123 fluorescence. It was established that the extent of mitochondrial membrane depolarization depends on the concentration of ACh applied to leech neurons (10^−6^ M Ach to 10^−2^ M ACh (Fig. 4B). The increase of ACh concentration in the bathing solution caused the elevation of Rh123 fluorescence intensity that reflects the depolarization of mitochondrial membrane. The dependence of the ΔΨamplitude of the mitochondria inner membrane during the Retzius cell AChR activation may be related to the Ca^2+^-influx during the AChRs activation [23]. It was found that two consecutive applications of ACh cause the reversible increase of Rh123 fluorescence intensity, which reflects the inner mitochondrial membrane depolarization. The second application of the ACh immediately after the restoration of fluorescence signal intensity led to the smaller response and longer recovery to the initial level of fluorescence. The observed effect is obviously due to the desensitization of the AChR [24].

The mitochondrial potential changes during RA are associated with the Ca^2+^ accumulation [25]. In this regard, the changes of inner mitochondria membrane potential in Retzius neuron during AChR activation was investigated upon reduction of extracellular Ca^2+^concentration. The absence of the extracellular Ca^2+^ does not completely block the development of the inner mitochondrial membrane depolarization during AChR activation. Our results suggest that the mitochondria depolarization is depended on Ca^2+^-influx from the extracellular medium and occurs, partly, due to the Ca^2+^-release from the intracellular compartments or plasma membrane desorption.

## 4. Discussion

It is known that in the Retzius cell, the localization of the clusters of vesicles with serotonin is associated with sites of the plasma membrane maximal depolarizations and electrostimulation caused a heterogeneous vesicles distribution in the different plasma membrane region [7]. So, the release of the vesicles with serotonin does not occur evenly throughout the neuron membrane, but there are areas for the formation of clusters of vesicles with serotonin and these areas correspond to the areas with the membrane potential local changes. Possible, the maximum OPD changes, localized in the near-membrane region of the Retzius neuron, where maximal changes in the membrane potential are observed and reflect the accumulation and the vesicles redistribution during AChR activation (Fig. 2A). Possible, that without AChR activation, the vesicle clusters and mitochondria are distant from the plasma membrane. The AChR activation evokes Ca^2+^ entry through L-type channels or membrane binding Ca^2+^desorption (Fig. 4). The last triggers exocytosis from vesicles that rest close to the plasma membrane and in parallel, activates ryanodine receptors and Ca^2+^-induced Ca^2+^ release, from endoplasmic reticulum. It is likely that the Ca^2+^ desorption leads to the neuron plasma membrane viscosity changes. The fast Ca^2+^ transient (Ca^2+^-channels or membrane binding Ca^2+^ desorption) reaches mitochondria, which respond by changes electron transport chain (ETC) and ATP synthesis. This activates the kinesin motors and vesicle clusters are transported towards the plasma membrane. The peripheral vesicle clusters and mitochondria receive more Ca^2+^ and ATP than the central clusters. As vesicles arrive at the plasma membrane and fuse, the 5-HT that had been released activates 5-HT2 receptors. These receptors are coupled to phospholipase C, which induces the formation of IP3, which activates Ca^2+^ release from submembrane endoplasmic reticulum and membrane fluidity changes. Ca^2+^ evokes further exocytosis, thus closing the local feedback loop.

## Author Contributions

Conceptualization, A.A., G.M.; methodology, O.S., A.I., A.A., T.K.; formal analysis, O.S., T.K., A.I.; investigation, A.I., N.K., A.A., G.M.; sources, O.S., A.I.; data curation, A.A., O.S., A.I.; writing—original draft preparation, G.M., T.K.; writing—review and editing, G.M., A.A., T.K. visualization, A.A., G.M.; supervision, G.M.; project administration, G.M.; funding acquisition, G.M. All authors have read and agreed to the published version of the manuscript.

